# Noise Invariance in Inferior Colliculus Neurons is Dependant on the Input Noisy Conditions

**DOI:** 10.1101/2020.03.03.975060

**Authors:** Maryam Hosseini, Gerardo Rodriguez, Hongsun Guo, Hubert H. Lim, Éric Plourde

**Author notes:** Corresponding author: Éric Plourde, NECOTIS, Department of Electrical and Computer Engineering, Université de Sherbrooke, 2500 Boulevard de l’Université, Sherbrooke, QC, J1K 2R1,Canada.

## Abstract

The auditory system is extremely efficient to extract audio information in the presence of background noise. However, the neural mechanisms related to this efficiency is still greatly misunderstood, especially in the inferior colliculus (IC). In fact, while noise processing under different conditions has been investigated at the auditory cortex level, studies in the IC have been much limited. One interesting observation has been that there seems to be some degree of noise invariance in the IC in the presence of white noise. We wish to broaden this knowledge by investigating if there is a difference in the activity of neurons in the IC, when presenting noisy vocalisations with different types of noises, input signal-to-noise ratios (SNR) and signal levels. We do so using a generalized linear model (GLM), which gives us the ability to study the neural activity under these different conditions at a per neuron level. We found that non-stationary noise is the only noise type that clearly contributes to the neural activity in the IC, regardless of the SNR, input level or vocalisation type. However, when presenting white or natural stationary noises, a great diversity of responses was observed for the different conditions, where the activity of some neurons was affected by the presence of noise and the activity of others was not. Therefore, there seems to be some level of background noise invariance as early as the IC level, as reported before, however, this invariance seems to be highly dependent on the noisy conditions.

**New & Noteworthy:** The neural mechanisms of auditory perception in the presence of background noise are still not well understood, especially in the IC. We studied neural activity in the IC when presenting noisy vocalisations using different background noise types, SNRs and input sound levels. We observed that only the non-stationary noise type clearly contributes to the neural activity in the IC. The noise invariance previously observed in the IC thus seems dependent on the noisy conditions.

## Introduction

Humans and animals have an amazing ability to separate relevant sounds from background noises. This ability is particularly important as a survival tool since it helps to gain an accurate understanding of the surrounding world. Even though perception in noise is an important part of auditory processing, the underlying neural mechanisms are still not well understood.

One major idea hypothesized to play a role in noise processing is adaptive processing, i.e. the ability of neurons to adapt their processing to the changes in the input stimulus statistics. It has long been proven that neurons in the auditory nerve (Wen et al. 2009; Robinson and McAlpine 2009; Zilany and Carney 2010; Wen et al. 2012), midbrain (Dean et al. 2005, 2008; Robinson and McAlpine 2009; Willmore et al. 2016) and cortex (Baccus 2006; Nagel and Doupe 2006; Watkins and Barbour 2008; Billimoria et al. 2008; Robinson and McAlpine 2009; Rabinowitz et al. 2011, 2012; Cooke et al. 2018) have the ability to adapt their processing to sounds with different intensities using processes called mean and contrast adaptation. What has newly been studied is the fact that neurons in different structures of the auditory pathway use mean and contrast adaptation to process the additive audio noise (Rabinowitz et al. 2013; Ding and Simon 2013). Ding and Simon 2013 studied contrast adaptation in the auditory cortex, as well as adaptive neural encoding of temporal modulations. They found that the very slow rhythm of speech (< 4 Hz) is robust to background noise. Rabinowitz et al. 2013 studied adaptation to stimulus statistics, mean and contrast, in different structures of the ascending auditory pathway and showed that the adaptation increases along the ascending auditory pathway. They also found that the structures that have the strongest adaptation, show the most noise invariance. In particular, the auditory cortex was shown to be more noise invariant than the inferior colliculus (IC), which in turn was shown to be more noise invariant that the auditory nerve. Therefore, neurons in the IC should respond to the addition of background noise to some degree. However, what remains to be answered is whether this neural response is different for different signal-to-noise ratios (SNR) or even different sound levels. Moreover, would different types of noises, e.g. stationary vs. non-stationary, affect differently this neural response?

Advanced processing such as predictive coding could play a role in generating different neural responses for different noise types. Predictive coding is the principle by which the brain tries to predict what will be the next input in a given sensory system and where any deviance from this prediction generates a stronger neural activity. It is known that predictive coding occurs in the cortex and it has been found that it increases along the auditory pathway. Therefore, the IC is less subject to predictive coding than the auditory cortex (Parras et al. 2017). Since non-stationary noise is by definition not predictable, it should generate more activity than stationary noise in the auditory cortex. It is however not clear if it will generate more neural activity than stationary noise in the IC as well.

In fact, most studies of the effects of noise on the activity of neurons in the auditory system have specifically focused on the auditory cortex (Schneider and Woolley 2013; Ding and Simon 2013; Moore et al. 2013; Mesgarani et al. 2014; Ni et al. 2016). One of the very few studies that have investigated noise processing in the IC is the Rabinowitz et al. 2013 study mentioned previously where the noisy conditions where limited to one stationary noise and one input sound level.

In this work we wish to study if there is a difference in neural activity in the IC when presenting different types of noises, and, moreover, when using different input SNRs and signal levels. In particular, we wish to investigate if the noise invariance reported for the neurons in the central nucleus of the IC (ICC) is present for all the studied conditions. To do so, we combine previously proposed metrics (Hosseini et al. 2019) that allow a per neuron interpretation, with recordings of neural activities in response to different types of background noises (non-stationary, white and natural stationary noises), SNRs (15 and 5 dB) and levels (55 and 65 dB). We found that non-stationary noise is the only one that clearly contributes to the neural activity in the ICC, regardless of the SNR, input level or vocalisation type. Moreover, we also observed that most ICC neurons respond to the non-stationary noise by a similar increased spiking activity. However, when we presented white or natural stationary noises, we observed a great diversity of responses where the activity of some neurons was affected by the presence of noise and others were not.

## Materials and Methods

### Anesthesia and surgery

Recordings were obtained from three guinea pigs. Animals were initially anesthetized with an intramuscular injection of ketamine (40 mg/kg) and xylazine (10 mg/kg). During the experiment, the anesthetic state was kept with supplements of ketamine and xylazine. It should be mentioned that using anesthesia affects neural responses to some degree, including the response latencies, the number of spontaneously active neurons and the threshold (Astl et al. 1996). To keep the body temperature at a natural state, a warm water blanket was used and the temperature was monitored with a rectal thermometer. The heart rate and oxygen concentration were also monitored. All experiments were carried out in accordance with policies of the University of Minnesota Committee on the Care and Use of Laboratory Animals.

Methods for surgeries, acoustic stimulation and neural recordings are similar to previous works and detailed in Lim and Anderson 2006. Briefly, the animals were fixed into a stereotaxic frame and the speaker was connected to the left ear through an ear bar inserted into the inner ear. Animals were given local anesthesia before opening the skull. The right occipital lobe was exposed and a multi-site electrode array was inserted in the ICC along the tonotopic axis. The ground wire was placed in the neck muscle. Probes (NeuroNexus, Ann Arbor, MI) with two shanks having 16 sites on each shank were used. Saline was added on top of the brain to prevent swelling. The experiments continued for 24 hours, which was the protocol limits. Three to four placements were obtained from each animal.

### Placement of electrode array in ICC

The desired placement of the electrodes were along the tonotopic axis of the ICC, as shown in Fig. 1. To find the correct positioning of the electrodes, we recorded the responses of neurons to different levels of broadband noise and pure tones. We then plotted the post-stimulus-time-histogram (PSTH) and frequency response maps (FRMs) of the neural responses to the noise and tones online to find the positioning of the electrodes in the ICC (Lim and Anderson 2006).

**Figure 1:**
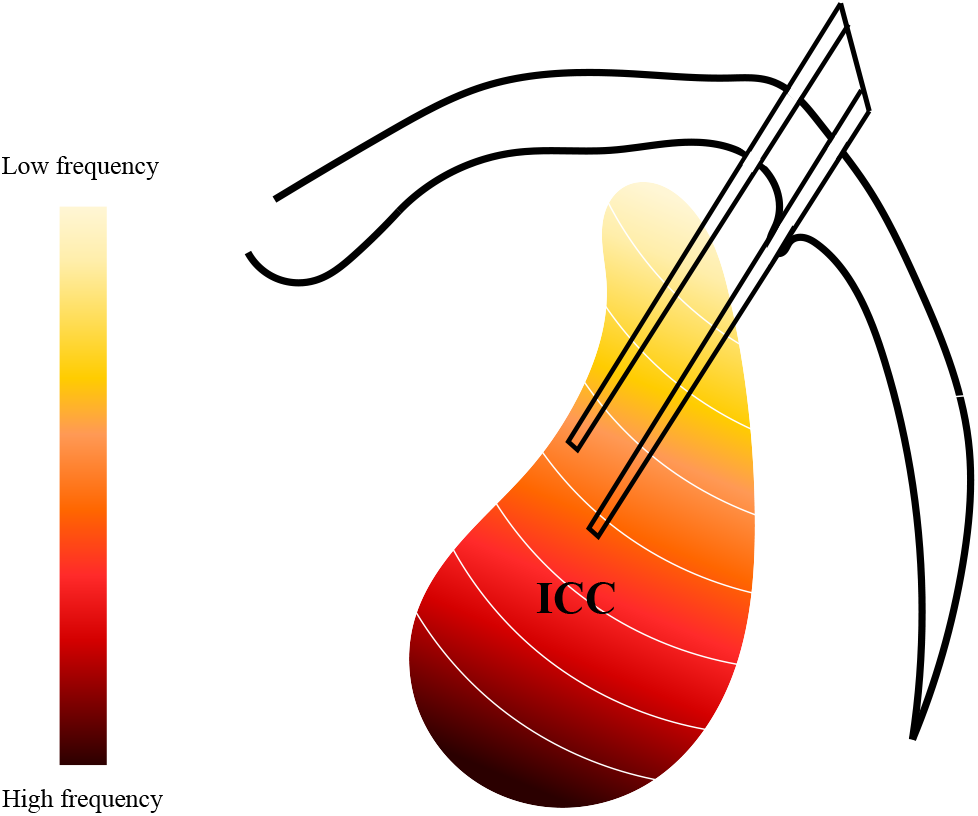
Placement of electrodes. The ICC has a laminar structure where the neurons on each lamina have a specific characteristic frequency (CF). The electrodes are implanted perpendicular to the iso-frequency laminas.

For the FRMs, pure tones had a length of 50 ms with 5 ms ramps at onset and offset, with levels ranging from 0 to 70 dB SPL with 10 dB steps. The frequencies of these tones varied from 1 kHz to 40 kHz at 8 frequencies/steps per octave. The broadband noise used for positioning the electrodes had a 50 ms duration with 0.5 ms ramps on the onset and the offset, with a level of 70 dB SPL. The frequency range of this broadband noises ranged between 0.625 kHz to 40 kHz, with a bandwidth of 6 octave.

### Acoustic stimulation and recording

Experiments were performed in a room that was acoustically and electrically insulated and were controlled from a computer located outside the room and connected to an acquisition system (TDT, Alachua, FL) using a custom software. Species-specific vocalisations (Rode et al. 2013) embedded in noise were presented to the guinea pigs and the multi-unit spiking activity of ICC neurons was recorded simultaneously for 32 sites, for 40 trials. Even though the activity recorded was a multi-unit activity, it has been shown that neighboring neurons in the ICC have similar characteristic frequencies (CF) and correlated firing and thus similar spectral and temporal preferences. In fact, the spectro-temporal receptive fields (STRFs) of neighboring neurons in the ICC show a strong similarity in CF and a moderately similar bandwidth, temporal response type, best modulation frequency, nonlinearity structure, and degree of information processing (Chen et al. 2012; Atencio et al. 2016). Interstimulus intervals were 476 msec. Each stimuls was randomly presented for each trial to correct for the effect of adaptation. We used three different noise types: non-stationary, white and natural stationary (i.e. wind) noises. These noises were added to three guinea pig vocalisations, named scream, squeal and tooth chutter, to obtain an SNR of 5 and 15 dB. The different noisy vocalisations were presented to the animals at a sound level of 55 and 65 dB. The temporal waveform and spectrogram of each vocalisation and noise can be seen in Fig. 2a and b respectively. Fig. 2c shows the spectrogram of the noisy vocalisations with different noises and at 5 dB SNR at a level of 55 dB.

**Figure 2:**
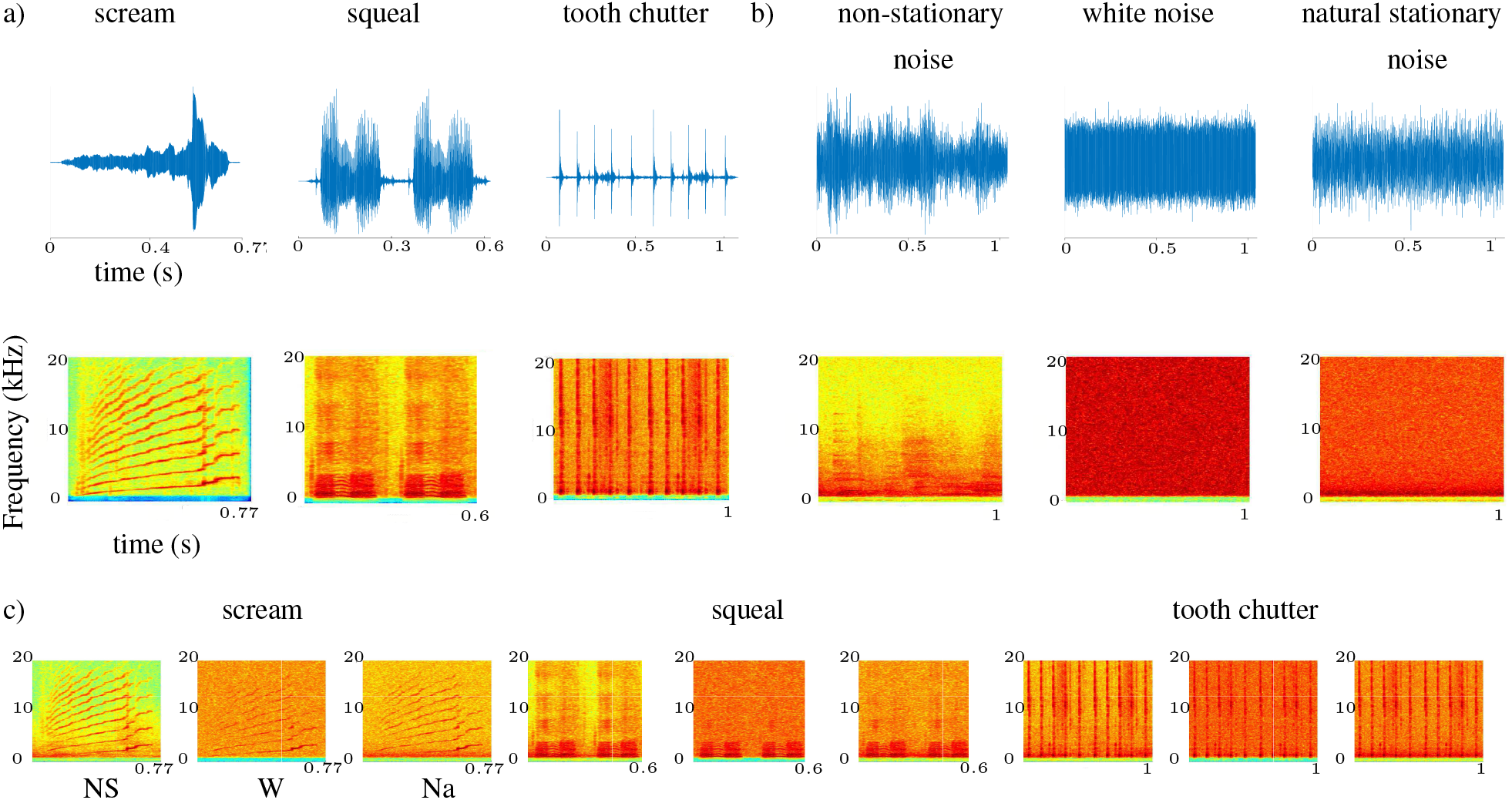
Acoustic stimuli used in the experiments: a) temporal waveform of the pure vocalisations (top) and their spectrogram representation (bottom), b) temporal waveform of the pure noises (top) and their spectrogram representation (bottom) and c) example spectrograms of noisy vocalisations at an SNR of 5 dB and an input level of 55 dB.

The same noises were used for all trials, placements and animals. The level of the pure vocalisations were held constant and the level of noise was altered to reach the desired SNR. The vocalization, noise and noisy vocalisation level at each SNR and level is displayed in Table 1. The sound level was calculated with the active level of the vocalisation, based on the International Telecommunication Union, Telecommunication Standardization Sector (ITU-T) recommendation P.56 (Kabal 1999; ITU-T 1993). The sampling frequency of the acoustic stimulation was 195.3125 kHz and the sampling frequency of the neural recordings was 24.414 kHz. Neural recordings started 60 ms before the presentation of the stimuli. Before being presented to the guinea pigs, the stimuli (clean vocalisation, pure noise and noisy vocalisations) were filtered between 1 kHz - 20 kHz using a bandpass filter to match the flat frequency band of the microphone used.

**Table 1:**
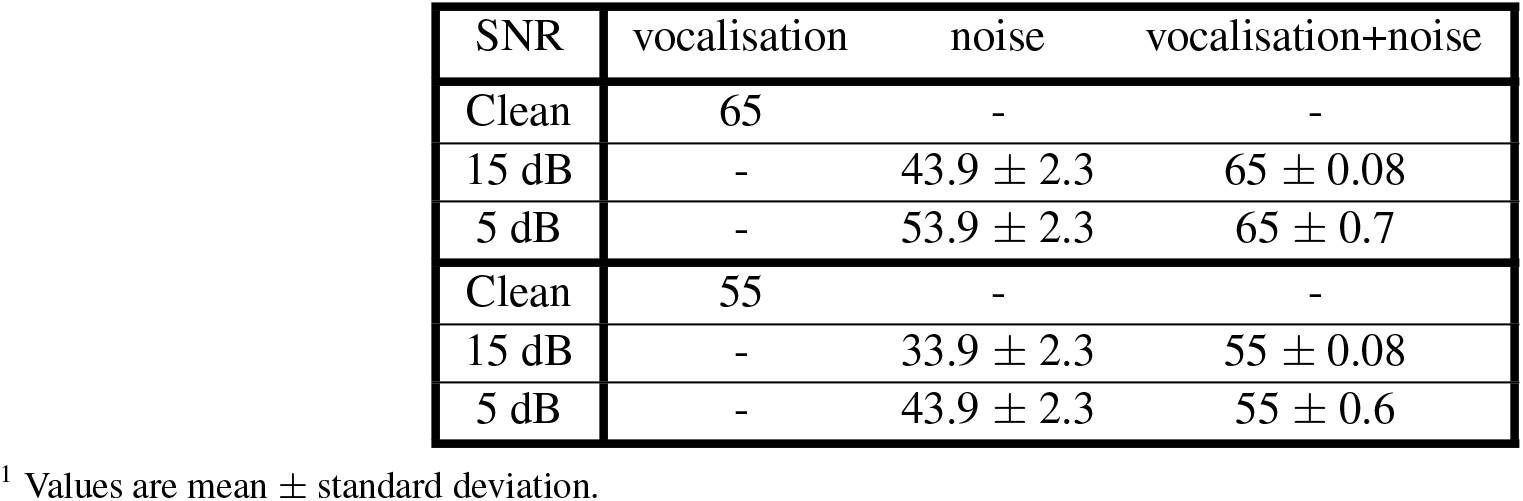
Noise and vocalisation sound pressure for the 65 dB and 55 dB input levels

### Sound calibration

Vocalisations, noises, noisy vocalisations, pure tones and broadband noise were presented to the animal through a speaker connected to the left ear through an ear bar. When these sounds pass through the speaker-earbar-ear canal system, their magnitude and phase change. To correct for these changes, we need to calibrate the sounds. For the case of pure tones and broadband noise, the recorded sound needed to be corrected only for magnitude. However, the vocalisations recorded by the microphone needed to be corrected also for phase. The procedure to calibrate the sounds was as described in Rode et al. 2013. In short, this was done by connecting the speaker to the earbar and then to a plastic tube that was used to replicate the guinea pig’s ear canal. The other end of the plastic tube was connected to a microphone. The sounds recorded at this end were used for calibration. To do that, we needed to find the inverse transfer function of the speaker-ear bar-ear canal system through the application of a normalized least mean squares (NLMS) adaptive filter.

### Offline data analysis and statistical analysis

The multi-unit activity recorded was filtered between 300 Hz and 3 kHz and a threshold of 3.5 times the standard deviation of the background noise was used to detect the spikes. The negative peaks were used to detect the spikes. The approximate CF for each site was found using FRMs and a custom GUI.

Only neurons that were responsive to the vocalisations were taken into account. To find these neurons, the baseline spiking rate of each neuron was calculated. Only the ones that had spiking rates that were one standard deviation (SD) higher than the baseline spiking rate in response to pure vocalisations were considered active. Of the total of 288 neurons recorded, 131 neurons for scream, 117 neurons for squeal and 158 neurons for tooth chutter were active.

All analysis were performed in MATLAB (Mathworks, Natick, MA). Interquartile range (IQR) was used as a measure of variability. Correlation between data points was studied using Pearson correlation coefficient and *p* values of two sample t-tests were used to compare data. Histograms, box plots and scatter plots were used to visualize the data.

### Neural activity metrics

One way to study the effects of noise on the spiking activity of neurons is to use PSTH based methods (Schneider and Woolley 2013; Moore et al. 2013; Ni et al. 2016). These methods assume linear-time-invariant (LTI) properties for sensory systems. However, sensory systems cannot be modelled as LTI (Theunissen et al. 2000; Escabı and Schreiner 2002; Lyzwa et al. 2016). A linear system is a system in which the output for a linear combination of inputs is the same as a linear combination of responses to those inputs presented individually and a time-invariant system is one where the output does not depend on the timing of the input. An LTI system has both properties. Since sensory systems are not LTI, the neural activity produced by the sum of the two stimuli will not be equivalent to the sum of the neural activity when presenting each stimulus separately. Moreover, another approach to model the effects of noise on the spiking activity of neurons is to try to reconstruct the input stimuli from the neural data (Mesgarani et al. 2014). However, such a reconstruction approach requires a great number of neurons to allow to draw conclusions and, furthermore, it is hard to conclude on the behaviour of specific neurons.

To avoid these problems, we developed metrics from a generalized linear model (GLM) that allowed us to study the effects of additive audio noise on the spiking activity of neurons. These metrics have previously been proposed in Hosseini et al. 2019.

Let us first describe the GLM, which will be followed by the two metrics used in this study. Point processes can be used to model the spiking activity of neurons. A point process can be completely described by its conditional intensity function (CIF) (Brown et al. 2003; Truccolo et al. 2005; Plourde et al. 2011). A general form for the CIF to model spiking activity is:

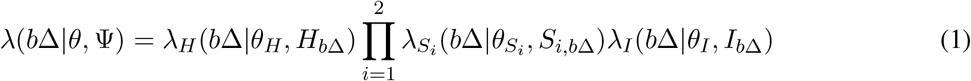

where the parameters to be estimated are 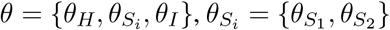 and where Ψ = {*H_bΔ_, S_i,bΔ_, I_bΔ_*} represents the known covariates with *b* ϵ {0, 1, …, *B* − 1} being the current time bin and Δ the bin size. The history of spiking activity is modelled by *λ_H_* (*b*Δ|*θ_H_, H_b__Δ_*), the effect of the vocalisation (*S*_1_) and noise (*S*_2_) on the neural activity is added to the model with 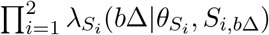 and *λ_I_* (*b*Δ|*θ_I_, I_b_*_Δ_) relates to the spontaneous activity of the neuron.

The *λ_H_*, 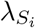 and *λ_I_* are generally modelled using exponentiated functions of the covariates. For *λ_H_*, modelling the history, we have:

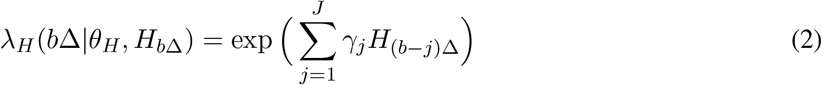

where *γ_j_* are the components of *θ_H_* and *H* is the spiking history.

The contribution of *i*th additive input, i.e. the vocalisation or noise, to the spiking activity is modelled as follows:

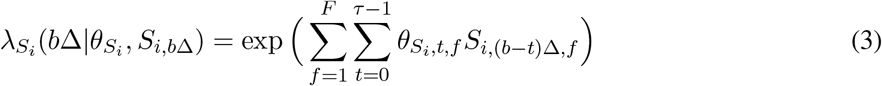

in which *S_i,bΔ,f_* is the representation of the vocalisation (*i* = 1) or noise (*i* = 2) at time *b*Δ and frequency band *f*.

The effect of the spontaneous activity on the spiking activity of neurons is modelled as:

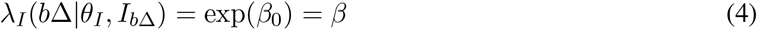

where the spontaneous activity is modelled as a constant term *β*_0_, which is to be estimated.

We can therefore rewrite (1) in the following form:

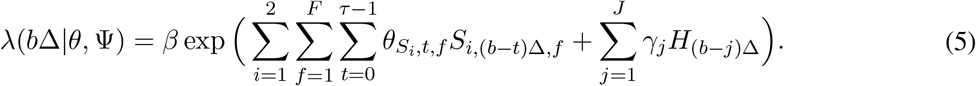

One way to interpret this GLM is that *λ_H_* and 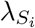 will modulate the spontaneous spiking activity *β*. For example, if a given input *S_i_* is zero, the value of 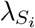 will be 1. This input will thus not modulate the spontaneous activity of the neuron. However, if a given 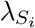 is greater or lower than 1, it will increase or decrease the spontaneous activity of the neuron and the corresponding input will thus contribute to the spiking activity of the neuron.

Two metrics have been derived from this model to study firstly, the influence of additive noise on the spiking activity of neurons and, secondly, the importance of the noise contribution to the neural activity compared to that of the vocalisations.

#### Metric no. 1: Influence of the additive noise on the neural activity

As mentioned previously, we can interpret *λ_H_* and 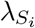 as modulators of the spontaneous firing rate. A measure of how the noise (*S*_2_) affects the baseline activity is therefore given by 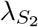 which we rename *F_noise_* here:

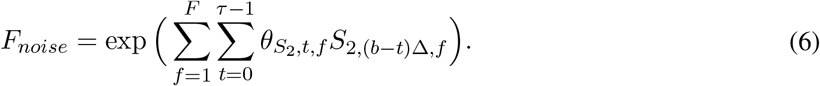

If the value obtained for *F_noise_* is higher or lower than 1, it will indicate that noise increases or decreases the firing activity respectively. A value close to 1 means noise has little to no effect on the spiking activity.

#### Metric no. 2: Importance of the noise contribution to neural activity compared to vocalisations

It would be relevant to have a measure of how important is the contribution of noise to the spiking activity of neurons compared to that of the vocalisation. We can define the modulation to the baseline activity caused by the vocalisation input, which we have named *F_voc_*, as follows:

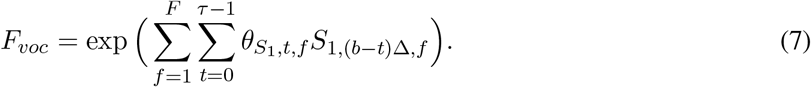

From this we define a metric, *F_ratio_*, that is the ratio of the contribution of noise over that of the vocalisation:

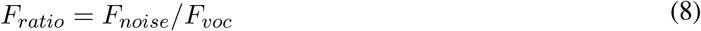

This metric is similar to a noise-to-signal ratio. We considered the noise-to-signal ratio instead of the more common signal-to-noise ratio because otherwise, a value of *F_noise_* ≃ 0 would yield a value of *F_ratio_* → ∞, while the proposed metric is mostly confined between 0 and 1.

We use these metrics to evaluate the effects of the noise on the spiking activity of ICC neurons. To do so, the GLM is fitted to the neural data using a truncated regularized iteratively reweighted least-squares (TR-IRLS) method (Komarek 2004; Plourde et al. 2011) where the time bin (Δ) is set to 1 ms.

Moreover, the stimuli are pre-processed as follows before being used in the GLM model. Firstly, the time domain input stimuli (clean vocalisations, pure noises and noisy vocalisations) are filtered between 1 kHz and 20 kHz. These band-limited stimuli are then processed with a gammatone filter to match the neuron’s CF, i.e. the frequency to which it best responds (Slaney 1993). Since neurons in the ICC are known to respond to the envelope of these bandpass signals (Rode et al. 2013; Woolley et al. 2006; Suta et al. 2003), the envelope of these bandpass signals are then extracted and used as input to the model for a given neuron.

### Goodness-of-fit of the GLM

To evaluate the goodness-of-fit of the GLM model to the recorded neural activity, we have used the time rescaling theorem (Brown et al. 2003; Haslinger et al. 2010). This theorem states that if the estimated CIF is a good approximation of the real CIF, then the rescaled times will be independent and uniformly distributed over [0,1). Kolmogorov Smirnov (KS) statistics was used to assess uniformity and the autocorrelation function (ACF) of the rescaled times was used to assess independence.

We used Kolmogorov-Smirnov (KS) plots of the distribution *F_m_* obtained from the model against the uniform distribution *F* to assess whether they lie within the 95 percent confidence bounds of the 45 degree line. We defined normalized KS statistics as (Plourde et al. 2011):

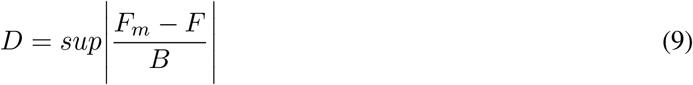

In which *B* is the width of the confidence bound. A good fit is achieved when *D* < 1, meaning the entire KS plot is within the confidence bound.

## Results

We now present the results regarding the effects of input background noise on the spiking activity of a given neuron in the ICC, when presenting noisy vocalisations. We first present examples of raster plots and PSTH for the spiking activity of a given representative neuron. In order to analyze the neural activity using the previously described metrics derived from the GLM, we then validate the goodness-of-fit of the GLM to the neural data. Results using the *F_noise_* and *F_ratio_* metrics, for three vocalisations (scream, squeal and tooth chutter), and three background noises (non-stationary, white and natural stationary noises) are then presented. We also study the effects of input sound level and SNR on the neural activity using levels of 55 dB and 65 dB with SNRs of 5 dB and 15 dB.

### Raster plots and PSTH

Fig. 3 shows examples of raster plots and PSTH of neural activity for clean scream, scream with non-stationary, white and natural stationary noises in the background and pure noises at a level of 55 dB and under 5 dB and 15 dB SNRs, for a given representative neuron with a CF of 2 kHz. Similar plots can be obtained for a level of 65 dB and other vocalisations.

**Figure 3:**
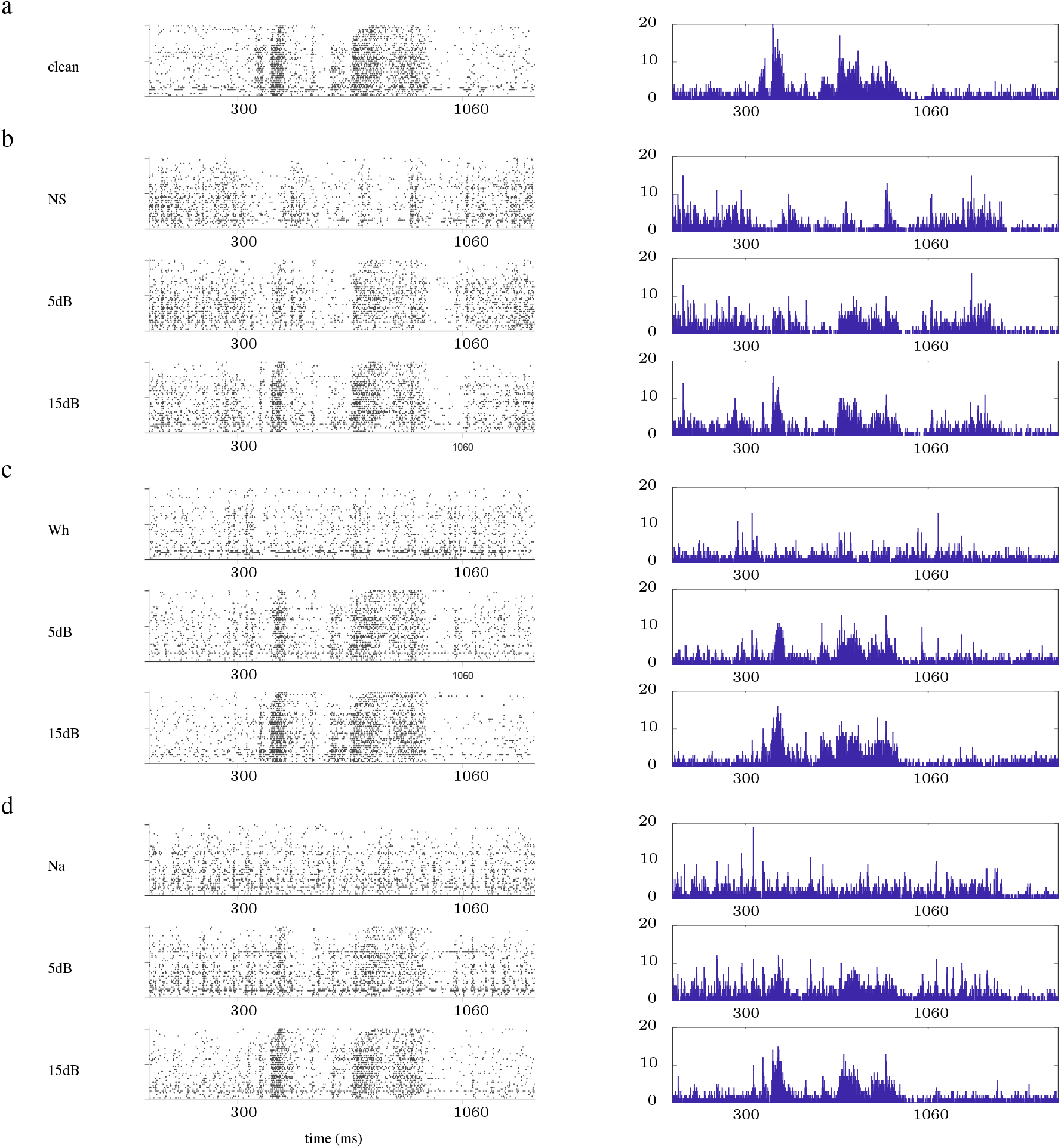
Raw data (raster (left) and PSTH (right) plots) for scream vocalisation with non-stationary (NS), white (Wh) and natural stationary (Na) background noises at a level of 55 dB. a) Plots for clean scream. b) Plots for a non-stationary (NS) noise in the background. The plots have been shown for pure noise (top), SNRs of 5 dB (middle) and 15 dB (bottom). c) Plots for a white noise (Wh) in the background. The plots have been shown for pure noise (top), SNRs of 5 dB (middle) and 15 dB (bottom). d) Plots for a natural stationary (Na) noise in the background. The plots have been shown for pure noise (top), SNRs of 5 dB (middle) and 15 dB (bottom). Similar results can be found for other vocalisations and a level of 65 dB. The neurons CF is 2 kHz

Apart from the fact that a lower SNR seems to produce more neural activity, it is rather hard to conclude anything using these representations.

### Goodness-of-fit of the GLM

Before being able to draw any conclusion from the GLM metrics, we need to verify if the GLM is a proper model for the neural data under consideration. Fig. 4 shows the box plots of normalized KS plots statistics, *D*, for different stimuli, noises, levels and SNRs. As indicated previously, a good fit is achieved when *D* < 1 It can be seen that the model fits extremely well the data for white and natural stationary noises, for all SNRs and levels. In fact, for all these cases, almost all of the normalized KS statistics values are below 1. While the fitting is slightly worse for the non-stationary noise, the model still fits the data very well since most of the normalized KS statistics values are below 1 or very close to it.

**Figure 4:**
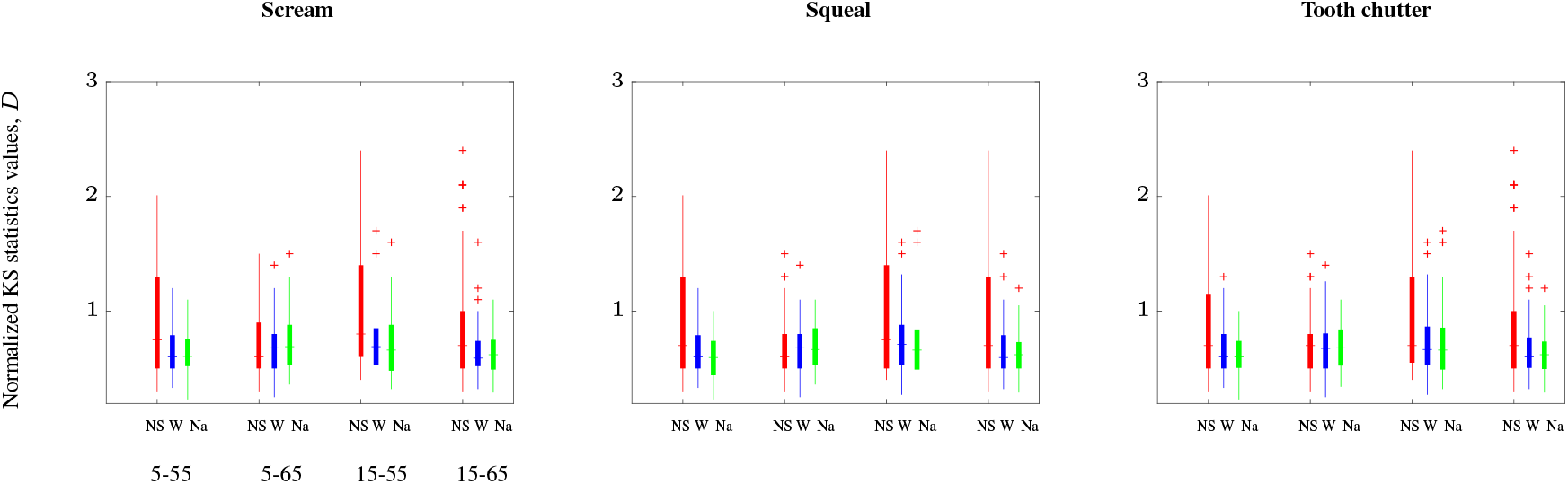
Box plot of normalized KS statistics values, *D*, for non-stationary noise (red, NS), white noise (blue, W) and natural stationary noise (green, Na), embedded in the background of vocalisations (scream (left), squeal (middle) and tooth chutter (right)). Each of the plots show the values for 55 dB and 5 dB SNR, 65 dB and 5 dB SNR, 55 dB and 15 dB SNR and 65 dB and 15 dB SNR. The plus signs (+) show the outliers of each case. The thick vertical line shows the middle 50 percent of the data and the whiskers show the 25th and 75th percentiles. The horizontal lines show the median of the data in each case. Since the *D* values are mostly below 1, it can be concluded that the rescaled times are uniform. Given the fact that the ACF are independent, we can thus conclude that the data fits the data very well for all SNRs and levels.

Using the ACF (results not shown), we observed that the majority of the rescaled times were highly independent for lags up to 100 ms (93.5% for scream, 90% for squeal and tooth chutter), therefore indicating that the rescaled times are independent.

The results from the KS plots and the ACFs therefore show that rescaled times are uniformly distributed and independent. The model thus is a great approximation to the neural activity and can be used for further analysis.

### Influence of the noise type on the neural activity

We study the effect of the noise type on the spiking activity of neurons in the ICC using the metric *F_noise_*. We have plotted the histograms of *F_noise_* values, for the different SNRs, noise and vocalisation types studied, with a level of 55 dB (Fig. 5) and 65 dB (Fig. 6) respectively. The red vertical line shows a value of *F_noise_* equal to 1, for which noise has no effect on the spiking activity of neurons.

**Figure 5:**
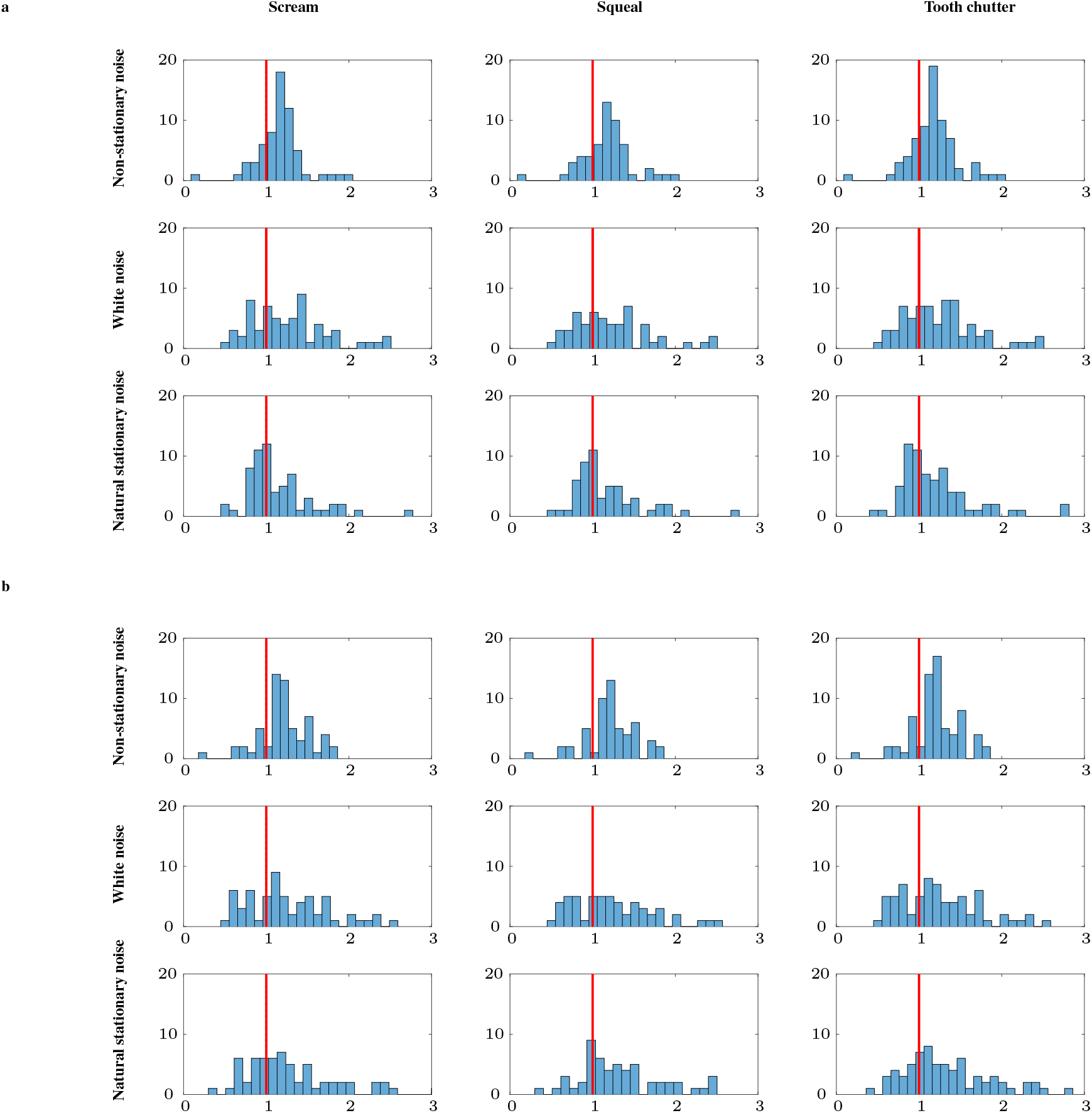
Distribution of the *F_noise_* values for different noises and vocalisations for a level of 55 dB and a) SNR of 5 dB and b) SNR of 15 dB. We present results for non-stationary, natural stationary and white noises as well as three vocalisations: scream, squeal and tooth chutter. From the responsive neurons, only the ones that had a good fit for all noises, SNRs and levels were considered (*n*=62 for scream, *n*=54 for squeal and *n*=69 for tooth chutter). The vertical red line corresponds to the *F_noise_* value of 1 indicating no contribution of the noise to the spiking activity. The presence of non-stationary noise clearly contributes to the spiking activity since the maximum of the distribution is always to the right of the red line. The presence of white noise at an SNR of 5 dB as well as the presence of white and natural stationary noises at an SNR of 15 dB do not seem to produce a unique neural signature where a wide variety of contribution to the neural activity can be observed. The presence of natural stationary noise at an SNR of 5 dB seems to indicate no contribution of the noise to the neural activity since the distribution is centered around the red line.

**Figure 6:**
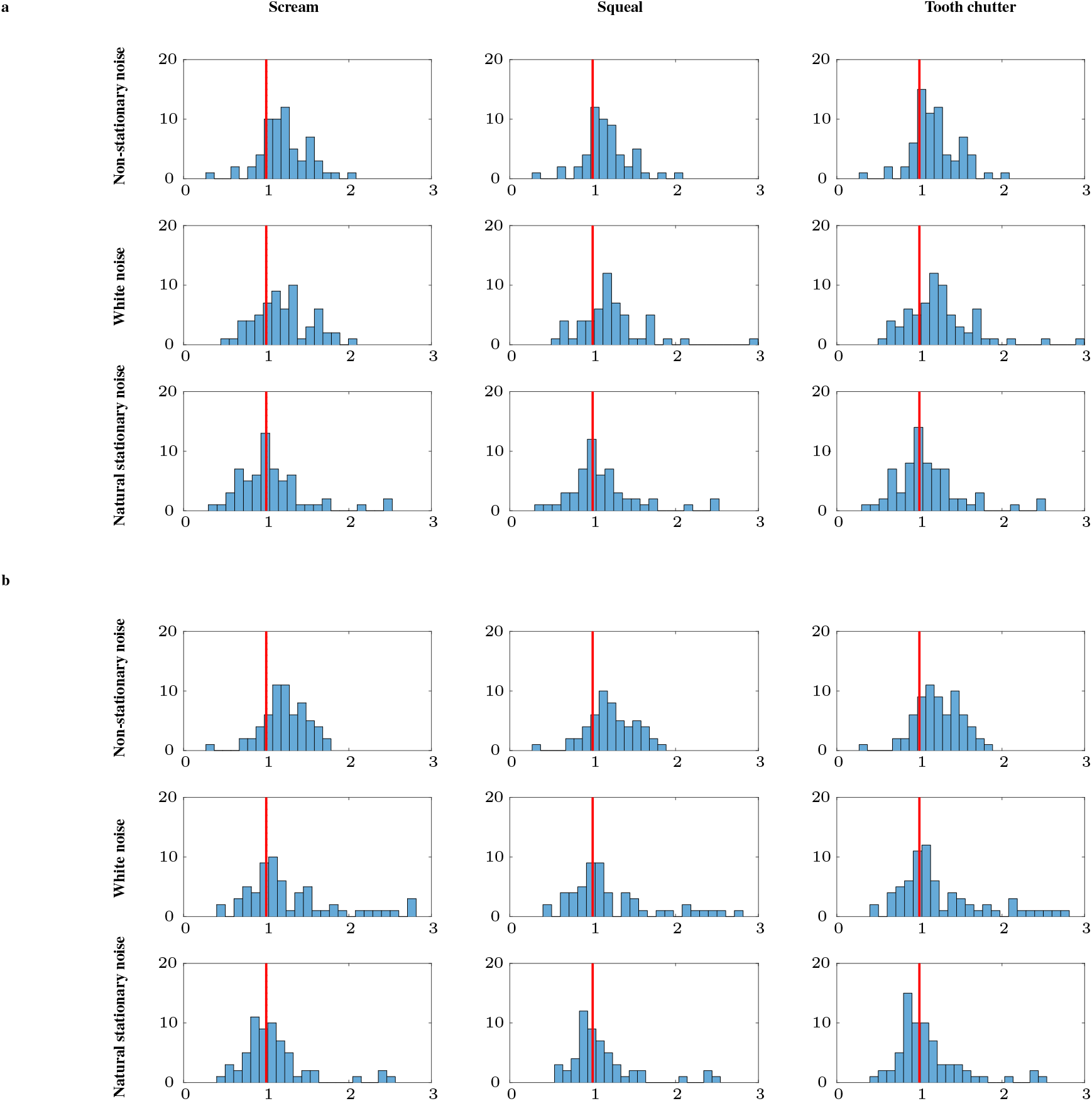
Distribution of the *F_noise_* values for different noises and vocalisations for a level of 65 dB and a) SNR of 5 dB and b) SNR of 15 dB. We present results for non-stationary, natural stationary and white noises as well as three vocalisations: scream, squeal and tooth chutter. From the responsive neurons, only the ones that had a good fit for all noises, SNRs and levels were considered (*n*=62 for scream, *n*=54 for squeal and *n*=69 for tooth chutter). The vertical red line corresponds to the *F_noise_* value of 1 indicating no contribution of the noise to the spiking activity. As for the 55 dB level results presented in Fig. 5, the presence of non-stationary noise clearly contributes to the spiking activity since the maximum of the distribution is always to the right of the red line. The presence of natural stationary noise at an SNR of 5 dB as well as the presence of white and natural stationary noises at an SNR of 15 dB seems to indicate no contribution of the noise to the neural activity since the distribution is centered around the red line. The presence of white noise at an SNR of 5 dB would seem to indicate a contribution of the noise to the neural activity.

As can be observed from Fig. 5 and 6, the distribution of *F_noise_* values seems to be quite different between different types of noises. In fact, the presence of non-stationary noise clearly contributes to the spiking activity since the maximum of the distribution is always to the right of the red line for all noise types, vocalisations, SNRs and input sound levels. With a wide variety of responses, the presence of white noise and natural stationary noises do not seem to produce a unique neural signature. In fact, for the 55 dB case, the distribution of neural responses seems flatter indicating a wide variety of responses, whereas for the 65 dB case, the majority of responses seem to indicate no contribution of the noises to the neural activity with a distribution centered around the red line.

To test the dispersion of values, we computed the IQR as shown in Table 2. A greater range indicates a wider dispersion of responses and therefore a greater diversity in neural responses indicating that neurons respond differently to the presence of a given background noise. From this table, it can be seen that generally, the IQR value is lower for non-stationary noise compared to the white or natural stationary noise types (except at 65 dB and 15 dB SNR), which shows also that in general, there is less variability in the values of *F_noise_* for non-stationary noise.

**Table 2:** Interquantile ranges for the scream, squeal and tooth chutter vocalisations corrupted with the non-stationary (NS), white (W) and natural stationary noises (Na) at the 55 dB and 65 dB input sound level and an SNR of 5 dB and 15 dB.

|  | Scream |  |  |  | Squeal |  |  |  | Tooth chatter |  |  |  |
| --- | --- | --- | --- | --- | --- | --- | --- | --- | --- | --- | --- | --- |
| Input level | 55 |  | 65 |  | 55 |  | 65 |  | 55 |  | 65 |  |
| SNR | 5 | 15 | 5 | 15 | 5 | 15 | 5 | 15 | 5 | 15 | 5 | 15 |
| NS | 0.26 | 0.34 | 0.33 | 0.35 | 0.27 | 0.29 | 0.24 | 0.37 | 0.26 | 0.28 | 0.27 | 0.39 |
| W | 0.53 | 0.72 | 0.39 | 0.55 | 0.51 | 0.69 | 0.34 | 0.54 | 0.5 | 0.7 | 0.36 | 0.57 |
| Na | 0.41 | 0.65 | 0.44 | 0.32 | 0.4 | 0.54 | 0.32 | 0.31 | 0.45 | 0.68 | 0.35 | 0.33 |

<sup>1</sup> IQR shows the dispersion of responses. The higher the IQR, the greater the diversity of the neural responses to the noises.
<sup>2</sup> Generally, we can observe lower values of IQR for non-stationary noise compared to white or natural stationary noises and 5 dB SNR compared to 15 dB SNR, indicating more similar spiking responses for different neurons in the former cases.

We can also observe that the IQRs for the SNR of 5 dB are generally smaller that that of 15 dB. This seems to indicate that the presence of a louder noise (5 dB SNR) produces more similar neural responses for different ICC neurons whereas the presence of a weaker noise (15 dB SNR) produce more diverse neural responses.

We also investigated if there was a similarity between the effect of different noise types on the spiking activity of a given ICC neuron. In other words, do two different noises produce the same spiking pattern for a given neuron. In order to do this, we have plotted in Fig. 7 the scatter plots of pairwise *F_noise_* values between two different noise types where we also show the *r* Pearson correlation coefficient. We present results here for the SNR of 5 dB and the 55 dB input level but similar results were obtained for other SNR and level combinations. The *r* Pearson correlation coefficient is very small, indicating that there is a weak correlation of the spiking activity when presenting two different types of noises to the same neuron. Therefore, the presence of a given noise type does not seem to have the same effect on the spiking activity of a neuron as the presence of another different type of noise.

**Figure 7:**
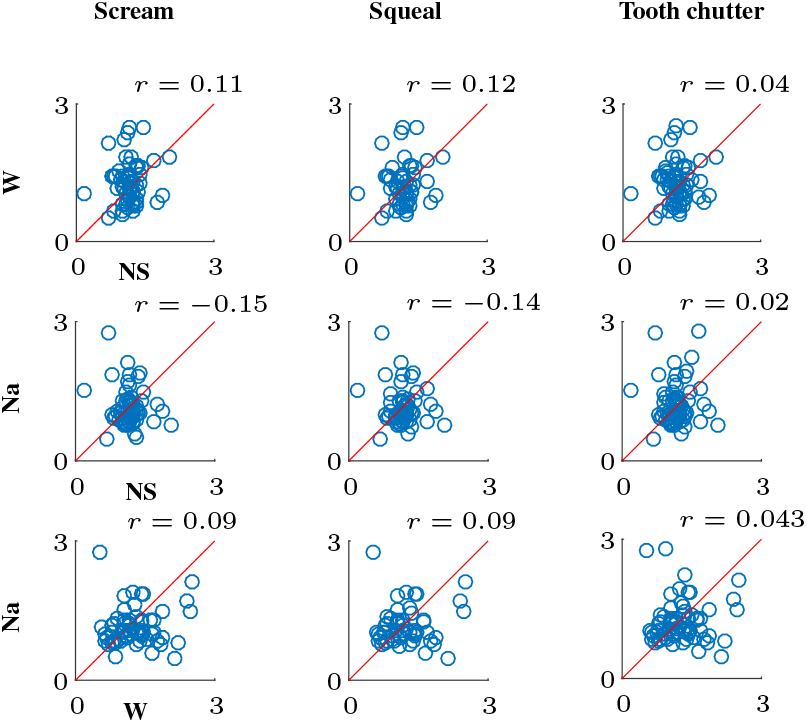
Scatter plot of *F_noise_* values for 55 dB and 5 dB SNR for paired noise types. Each point indicates the *F_noise_* values for non-stationary noise (NS), white noise (W) or natural stationary (Na) noise, as indicated by the axis names, for a given neuron. *r* is the Pearson correlation coefficient value between each pair. The *r* Pearson correlation coefficient is very small, indicating a weak correlation of the spiking activity when presenting two different types of noises to the same neuron. Therefore, different noise types don’t seem to affect the spiking activity of a given neuron in the same manner.

### Correlation between *F_noise_* and the characteristic frequency of neurons

In this section, we have investigated the correlation between the values of *F_noise_* and the CF of ICC neurons. Fig. 8 shows the scatter plot of *F_noise_* vs. the CFs for different vocalisations and noise types at an input sound level of 55 dB and an SNR of 5 dB. Similar results were obtained for other sound level/SNR combinations.

**Figure 8:**
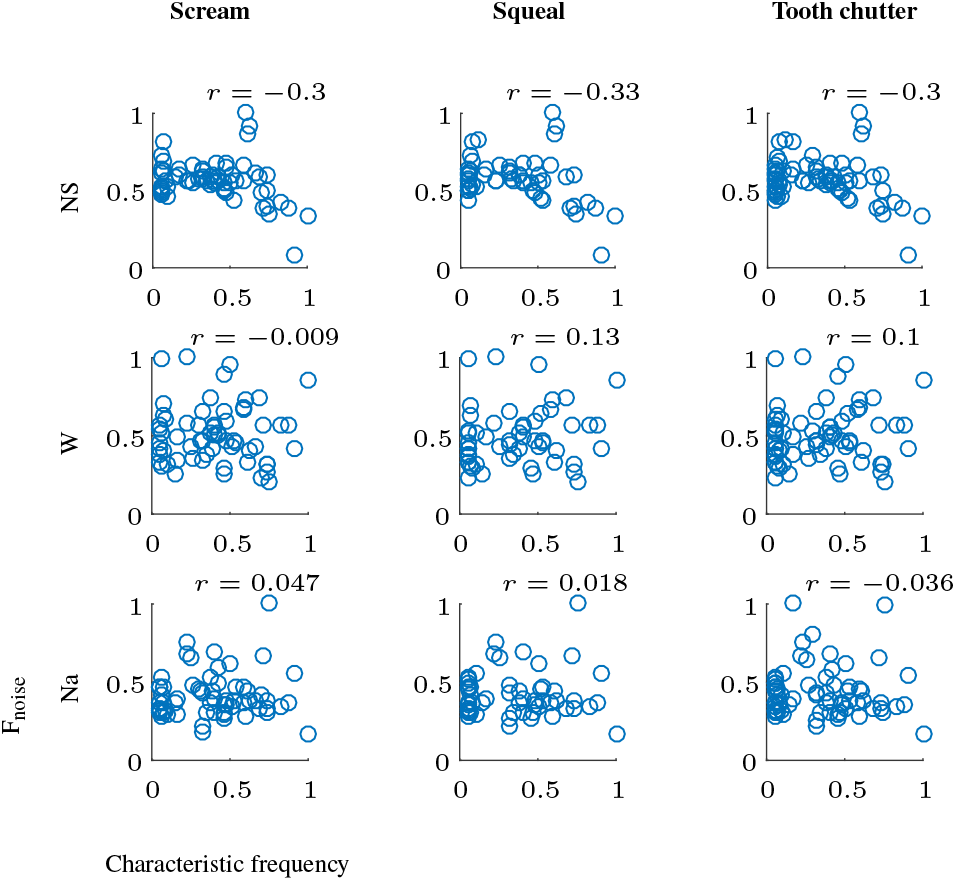
Scatter plot of the normalized *F_noise_* values vs. normalized characteristic frequencies of neurons for a 55 dB level and 5 dB SNR. Each point indicates the normalized *F_noise_* value vs. normalized characteristic frequency of a given neuron for non-stationary noise (NS), white noise (W) or natural stationary (Na) noise. *r* is the Pearson correlation coefficient value between each pair. A negative correlation exists between the CF and the contribution of the NS noise to the neural activity where lower CF neurons contribute more.

We observed a negative correlation between the effect of the noise on the neural activity and the CF of the neuron when presenting non-stationary noise. This indicates that the neural activity of lower CF neurons is more affected by the presence of noise than that of higher CF neurons. On the other hand, when presenting white or natural stationary noises, there doesn’t seem to be any relationship between CF and the values of *Fnoise*.

This could be caused by the fact that non-stationary noise has stronger components at lower frequencies, as can be seen from Fig. 2a. Since noise is not strong enough at higher frequencies it would generally cause less distortions. For white and natural stationary noises, all frequency components have the same power, which would cause them to affect all frequencies similarly.

### Effect of vocalisation type on the neural activity

Table 3 shows pairwise comparison between the values of *F_noise_* between two different vocalisations for the same noise and level. The *p* value shown is the smallest for different SNR and level combinations. The *p* values in the table are high (*p* > 0.37, two sample *t*-test). This shows that, in general, noises have similar effects on the spiking activity of neurons irrelevant of the vocalisation type.

**Table 3:**
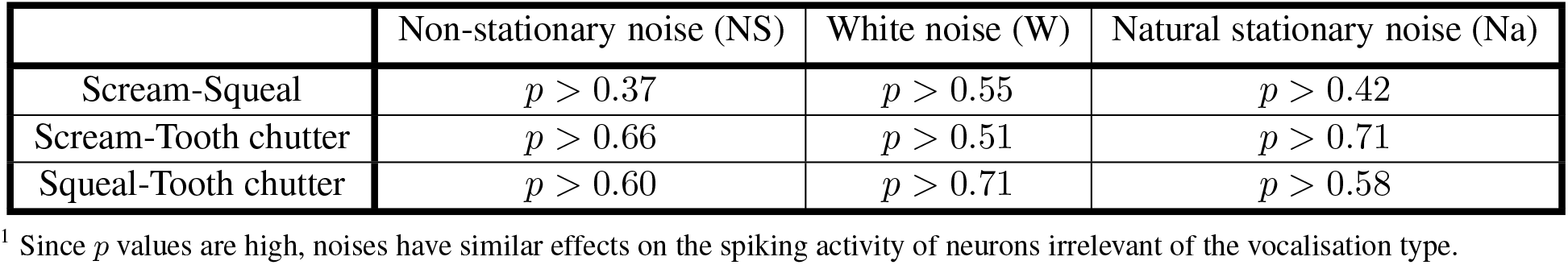
Two sample *t*-test *p* values for the pairwise comparison of *F_noise_* values between two different vocalisations for the same noise and input sound level.

|  | Non-stationary noise (NS) | White noise (W) | Natural stationary noise (Na) |
| --- | --- | --- | --- |
| Scream-Squeal | $p > 0.37$ | $p > 0.55$ | $p > 0.42$ |
| Scream-Tooth chatter | $p > 0.66$ | $p > 0.51$ | $p > 0.71$ |
| Squeal-Tooth chatter | $p > 0.60$ | $p > 0.71$ | $p > 0.58$ |

<sup>1</sup> Since $p$ values are high, noises have similar effects on the spiking activity of neurons irrelevant of the vocalisation type.

### Influence of noise on the neural activity compared to the vocalisation

This section investigates the contribution of the noise to the spiking activity of an ICC neuron compared to that of the vocalisation. To do that, we have plotted the box plot of the *F_ratio_* values in Fig. 9. From this plot it can be seen that, regardless of the type of noise, vocalisation or SNR, most of the values of *F_ratio_* are lower than 1, meaning that the effect of any type of noise on the spiking activity of neurons is always lower than the effect of the vocalisation. This shows that the main input driving the neural activity in the studied conditions is always the vocalisation. Even though at this stage of the auditory processing, noises are clearly still encoded in the neural activity, this homogeneity in the strong neural responses to the vocalisations proves that the processing of background noise and the segregation of the relevant auditory information starts as early as the ICC.

**Figure 9:**
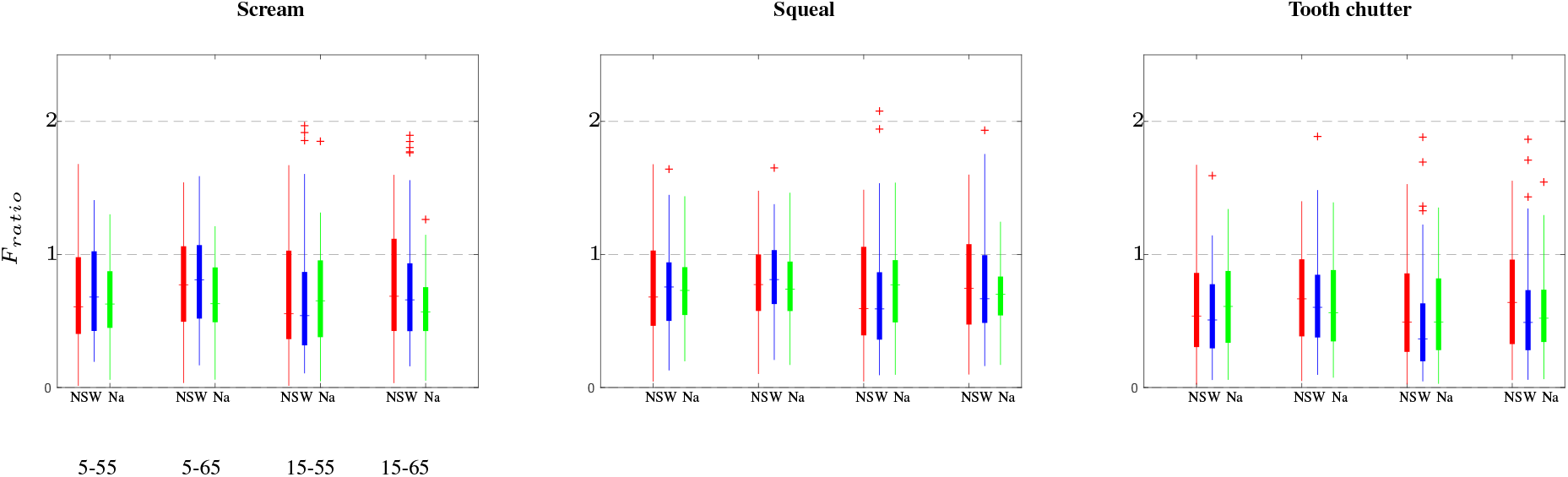
Box plot of *F_ratio_* values for non-stationary noise (red, NS), white noise (blue, W) and natural stationary noise (green, Na), embedded in the background of vocalisations (scream (left), squeal (middle) and tooth chutter (right)). Each of the plots show the cases for a 55 dB level and 5 dB SNR, 65 dB and 5 dB SNR, 55 dB and 15 dB SNR as well as 65 dB and 15 dB SNR. The plus signs + show the outliers in each case. The thick vertical line shows the middle 50 percent of the data values and the whiskers show the 25th and 75th percentiles. The short horizontal lines show the median of the data in each case. *F_ratio_* values are always lower than 1 indicating that, regardless of the type of noise, vocalisation or SNR, the vocalisation always has a greater effect on the neural activity than the background noise.

## Discussion

We have studied the effect of background noise on the neural activity of ICC neurons using different noise types at different SNRs and input sound levels, as well as different vocalisations. The effect of either a lower SNR or a higher input level, two conditions where the noise level is higher, seems to produce neural activity that is more similar amongst different ICC neurons than for either a higher SNR or lower input level, where the noise levels are weaker. This was more pronounced for white and natural stationary noises than for non-stationary noise. This would thus suggest that the adaptation to the mean of the stimulus used by neurons and observed in previous studies (Dean et al. 2005; Baccus 2006; Nagel and Doupe 2006; Watkins and Barbour 2008; Dean et al. 2008; Billimoria et al. 2008; Robinson and McAlpine 2009; Wen et al. 2009; Zilany and Carney 2010; Rabinowitz et al. 2011, 2012; Wen et al. 2012; Willmore et al. 2016; Cooke et al. 2018), i.e. that neurons adapt their neural activity to the input sound level, might not be true for all ICC neurons, especially in the presence of stationary noises. In fact, there might be groups of neurons in the ICC that respond to noises differently than other groups. In fact, this has already been shown to be the case in the auditory cortex (Ni et al. 2016).

Moreover, most of the values of *F_noise_* for natural stationary noise were concentrated around 1 which means that natural stationary noise almost doesn’t impact the spiking activity of neurons. This is to be expected since some neurons in the ICC are thought to respond to the envelope of an input sound and the envelope of a non-stationary noise is varying much more than the envelope of a stationary noise. Surprisingly, although white noise is stationary, it seems to have an impact on the neural activity for the specific case of a 65 dB level and 5 dB SNR, i.e. where the noise level is the highest of all the conditions studied. The reason could be due to a process called stimulus specific adaptation (Malmierca et al. 2009). In fact, white noise is an unfamiliar sound while a natural stationary sound, wind in our case, has possibly been heard before. The confirmation of this hypothesis would however necessitate further investigations.

Finally, we observed that the noise invariance reported in previous studies for ICC neurons (Rabinowitz et al. 2013; Ding and Simon 2013) seems to be dependent on the noise type. In fact, most of the previous studies used stationary noises and in fact, we observe that stationary noises (white and natural stationary) do seem to affect less the neural activity of ICC neurons. However, we observe that non-stationary noise greatly affects the neural activity of neurons in the ICC. It would remain to be investigated if noise invariance in the presence of non-stationary noise is also weaker in the auditory cortex.

## Acknowledgements

This work was supported by the Natural Sciences and Engineering Research Council of Canada (NSERC).

